# A systems approach to B-cell development identifies BDNF as a regulator of human B lymphopoiesis

**DOI:** 10.1101/2025.08.29.673055

**Authors:** Neta Nevo, Yuval Klein, Ayelet Alpert, Amir Grau, Doron Melamed, Timothy J. Few-Cooper, Neta Milman, Shai S. Shen-Orr

**Author notes:** These authors contributed equally (Timothy J. Few-Cooper, Neta Milman, Shai S. Shen-Orr). **Corresponding author**: Shai S. Shen-Orr, Department of Immunology, Rappaport Faculty of Medicine, Technion-Israel Institute of Technology, 1 Efron St., Haifa, 3525433, Israel.

## Abstract

B-cell aplasia is a major consequence of aging, chemotherapy, and B-cell-depleting immunotherapies, compromising immune protection against infections, cancer, and vaccines. Yet, unlike the myeloid and erythroid lineages, no strategy exists to accelerate human B-cell reconstitution. Here, we used an integrative systems biology approach to identify regulators of human B lymphopoiesis in the bone marrow (BM) microenvironment. By combining single-cell transcriptomic analysis of human BM with intercellular communication mapping, we generated an initial set of candidate factors predicted to act on developing B cells. To distinguish biologically meaningful putative regulators from a broad candidate space, we further intersected these findings with orthogonal human datasets capturing age-impaired B lymphopoiesis and protein dynamics associated with B-cell depletion and reconstitution. This convergent prioritization strategy highlighted a focused set of putative regulators, among which brain-derived neurotrophic factor (BDNF) emerged repeatedly as a top putative regulator. Functional interrogation in progenitor BM cells showed that several prioritized putative regulators induced transcriptional programs linked to early immune development, with BDNF consistently promoting pathways associated with B-cell differentiation. Importantly, in a human *in vitro* BM co-culture system, BDNF enhanced the differentiation of CD34+ hematopoietic progenitors into CD19+ progenitor B cells. Together, these findings identify BDNF as a previously unrecognized regulator of early human B lymphopoiesis and establish a general framework for uncovering functional hematopoietic regulators by integrating single-cell analysis with complementary biological and clinical signals. This approach may support future strategies to improve immune reconstitution in settings of prolonged B-cell depletion.

## Introduction

B-cell aplasia, defined as a marked decrease in B lymphocyte count, has long been recognized as a complication of both chemotherapy treatment and aging ^1–7^. More recently, however, the clinical burden of B-cell aplasia has intensified with the widespread use of B-cell targeted immunotherapies ^8–11^.

Notably, aging alone is associated with a skewed shift in bone marrow (BM) output toward the myeloid lineage at the expense of lymphoid differentiation reducing baseline levels of B cells in older individuals ^12–14^. Moreover, age-related impairments in B lymphocyte regeneration in the BM ^15,16^ and alterations in B-cell subset composition are well established ^17,18^.

As the cornerstone of humoral immunity ^19,20^, the lack of functional, mature B lymphocytes results in significant complications including severe infections notably with encapsulated bacteria ^21–23^, impaired tumour surveillance and decreasing the response to biological treatment ^24,25^. Yet, unlike the myeloid and erythroid lineages, whose constitution can be accelerated by G-CSF and Erythropoietin respectively, no BM B-cell growth factors are currently available in the clinic. Instead, patients with sustained B-cell depletion are treated with infusions of pooled immunoglobulins. However, such treatment is associated with immediate and severe side effects ^26^. Additionally, it does not reduce the incidence of infections ^27,28^. Therefore, identifying novel strategies to accelerate the development of B cells is of increasing clinical value.

B lymphocytes are derived from hematopoietic stem and progenitor cells (HSPC) in the BM. Through sequential, multistep developmental stages, HSPCs differentiate into progenitor B cells and subsequently, immature B cells that migrate toward peripheral blood (PB) and secondary lymph nodes for further maturation ^20^. Key regulators of B lymphopoiesis, C-X-C Motif Chemokine Ligand 12 (CXCL12) and Interleukin (IL) 7, are secreted from mesenchymal stromal cells (MSC) residing within cellular niches in the BM ^29,30^. However, such factors are not used to enhance the development of B cells in clinical settings due to their lack of specificity, and oncogenic potential ^31–36^. B lymphopoiesis is further supported by osteoblastic and Treg-mediated microenvironmental signals, and complex transcriptional and epigenetic networks which collectively act through a multi-step, tightly regulated process that is difficult to mimic^37–39^.

Recent technological advances have improved our ability to simultaneously measure numerous markers at the single-cell level. Indeed, previous studies have traced the process of B lymphopoiesis in the BM using single cell methods ^40–46^, yet most of knowledge in this field is based on works in rodents, which demonstrate significant differences in the composition and biological function of their BM B cells relative to humans ^47–50^. Investigation of human BM using single cell expression and epigenetic profiling further exposed the progenitor B-cell landscape ^51–54^. However, these achievements have not been translated into the clinic.

To discover regulators for B lymphopoiesis, we harnessed an integrative and systems-level approach, leveraging both clinically derived data and biological experiments. Through in-depth characterization of the BM B lymphopoiesis process, we identified, prioritized and validated candidates - toward accelerating the development of B cells in the BM. Notably, our approach revealed the potential for the neurotrophic factor, Brain-Derived Neurotrophic Factor (BDNF), to regulate human B-cell development, an unappreciated function not yet ascribed to this factor.

## Material and Methods

### Computational Methods

#### Mapping the human bone marrow B-cell landscape

To study human BM B-cell development, we analyzed a published scRNA-seq dataset (GSE149938) comprising 7,643 cells and 18,008 genes. Data were preprocessed, normalized, and dimensionally reduced to enable trajectory inference and transcriptomic visualization. Cell type annotations were validated against canonical B-lineage markers spanning the full developmental axis, from HSPCs through to mature plasma cells (see Supplemental Methods and Supporting Information S1: Figure 1A).

**Figure 1:**
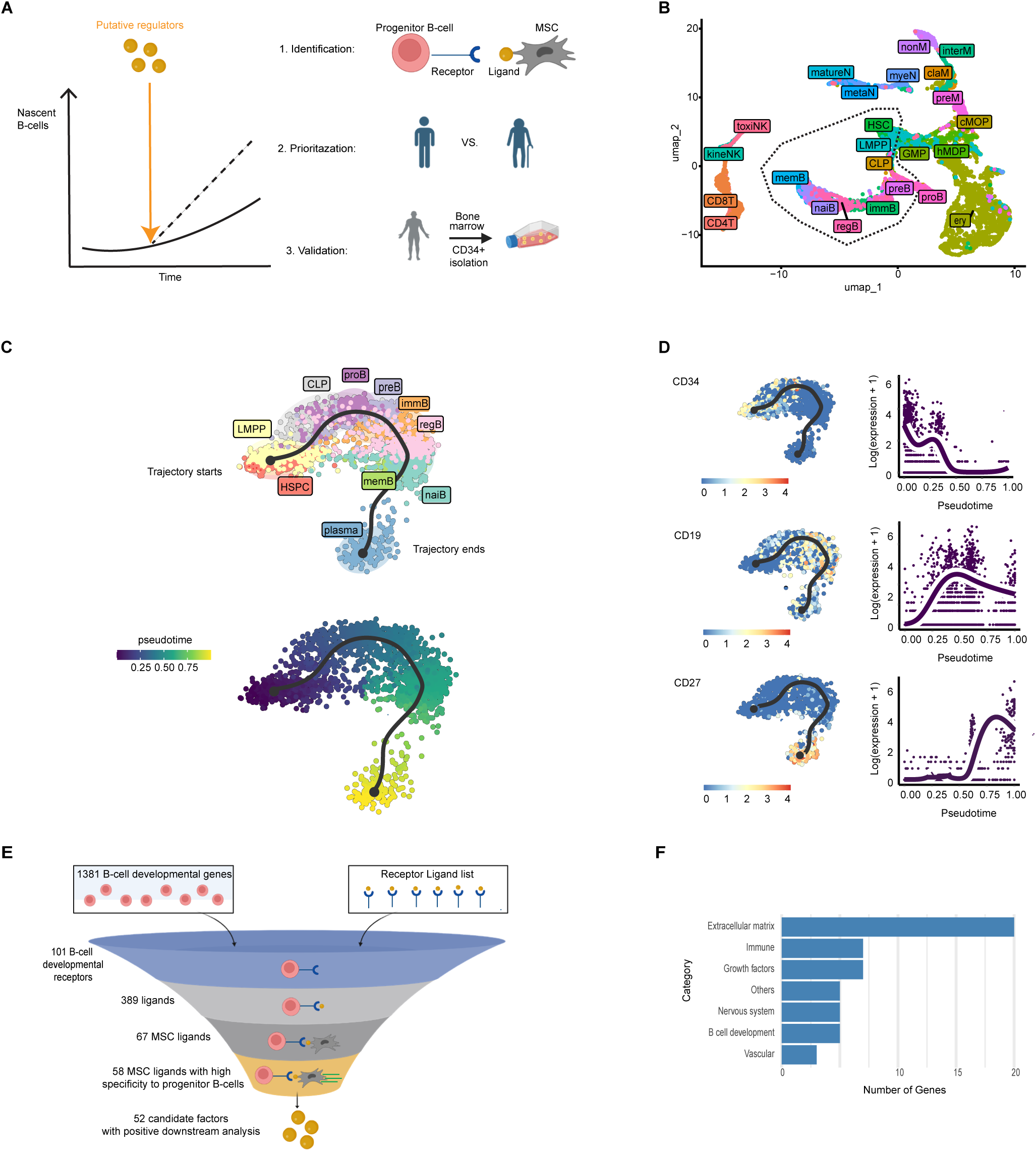
Mapping intercellular communication in human bone marrow identifies candidate factors predicted to influence B-cell development. **(A)** A schematic depicting the study workflow for identification and prioritization of putative B-cell regulators toward enhancing the development of B cells. The workflow consists of three major steps: (i) the identification of intercellular interactions in the BM; (ii) the prioritization by selection of factors playing a role in B-cell development upon aging; and (iii) validation of findings using *in-vitro,* biological systems. BM, Bone Marrow. **(B)** UMAP dimensionality reduction derived from analysis of publicly-available scRNA-seq data (GSE144938). 2127 progenitor B cells, marked by dashed lines, were utilized for inferring a pseudotime developmental trajectory where each dot represents a cell coloured based on high-resolution cell-typing. (**C**) Left - A trajectory of 2127 B-cell progenitors initiating at HSPCs through LMPP, CLP, proB, preB, immB, regB, memB, immB, naiB and plasma cells. Right - pseudotime scores ranging from 0 (HSPC) to 1 (Plasma cells) (right). **(D)** Scaled expression of three hallmark B-cell developmental genes: CD34, CD19 and CD27, as a function of pseudotime. For each marker, low and high expressions are denoted as blue and red respectively, and an associated graph displays the scaled expression pattern along the trajectory. **(E)** An illustration depicting our stepwise strategy to characterize and short-list putative B-cell regulators by integrating the genes of nascent B cells and MSCs with R-L pair lists. **(F)** A bar plot summarizing the distribution of significant biological pathways associated with the 52 identified putative B-cell regulators, as inferred from Gene Ontology annotations. MSC, Mesenchymal stromal cell; R:L, Receptor:Ligand; UMAP, Uniform Manifold Approximation and Projection; HSPC, Haematopoietic Stem and Progenitor cells; LMPP, ;CLP, Common Lymphoid Progenitors; immB, Immature B; regB, regulatory B; memB, memory B; immB, immature B; naiB, naive B.

#### Differential gene expression and trajectory inference

To identify genes dynamically regulated during B-cell development, we performed differential gene expression analysis, retaining genes meeting a corrected significance threshold. B-cell developmental trajectories were subsequently inferred using pseudotime modeling, and genes significantly associated with pseudotime were prioritized based on a Wald statistic threshold derived from the inflection point of the statistic’s distribution curve (Supporting Information S1: Figure 2). Applied sequentially, these filters reduced an initial set of 3,914 candidate genes to 1,381 trajectory-associated genes for downstream analysis. Full filtering parameters and software specifications are provided in the Supplemental Methods.

**Figure 2:**
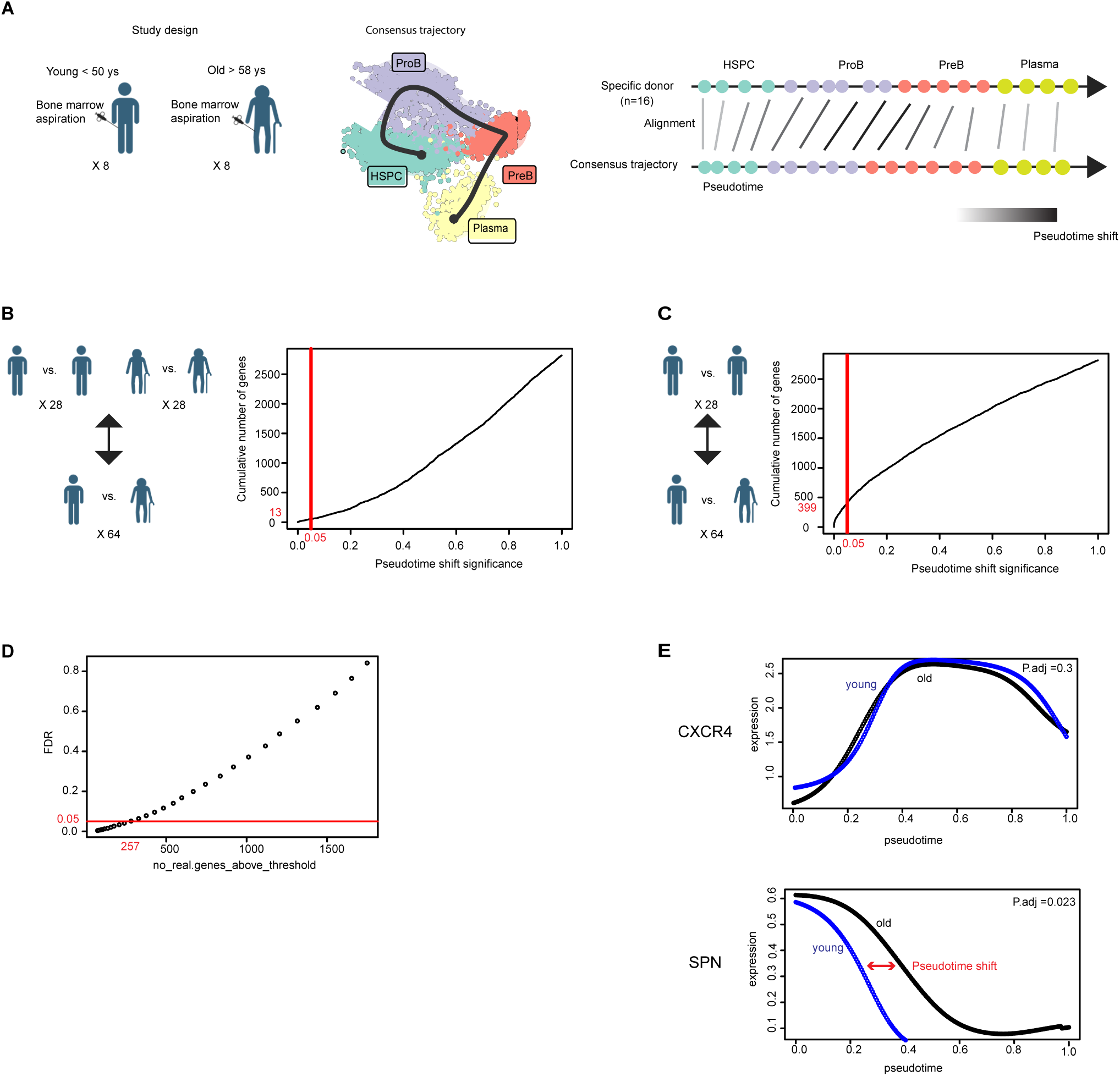
Pairwise trajectory alignment reveals age-dependent differences in the expression dynamics of pseudotime associated genes. **(A)** Left - A schematic depicting our approach to characterizing the differences in longitudinal, expression dynamics during B lymphopoiesis upon aging. A consensus B-cell developmental trajectory generated utilizing published, healthy human BM scRNA-seq data including 16 donors aged 24-67 revealed four major, expected subpopulations: HSPC, proB, preB and plasma cells. Right - The alignment procedure (via the *cellAlign* algorithm), as conducted between each time point of each donor and the consensus trajectory, captures pseudotime shift differences i.e. gaps in pseudotime on a per-gene basis and highlights age-related discrepancies in expression dynamics. **(B)** Illustrations and histograms showing comparisons of expression dynamics within the same age groups (28 pairwise comparisons within 8 young individuals vs. 28 pairwise comparisons within 8 old individuals) and across age (64 pairwise comparisons between 8 young individuals and 8 old individuals), which did not yield any significant results (p.adjust < 0.05). **(C)** As in (b), but confined to pairwise comparisons conducted solely within young individuals vs. older individuals, which identified 399 genes significant results (p.adjust < 0.05). **(D)** A scatter plot displaying FDR permutation tests aiming to identify the probability that any given pseudotime-shifted gene may reach significance by chance (Methods). The plot shows the number of genes (x-axis) with FDR greater than or equal to a given threshold (y-axis). Each point along the curve represents the cumulative count of genes that exceed the corresponding FDR value. The red dashed vertical line indicates a threshold of FDR = 0.05, below which 257 genes remained. **(E)** Line plots of two B-cell developmental receptor expression patterns across pseudotime without (CXCR4) and with (SPN) pseudotime shifts, as inferred from *cellAllign* analysis.

#### Receptor-ligand interaction mapping

To identify putative signals from BM MSCs capable of instructing B-cell development, we constructed an integrated receptor-ligand reference by harmonizing three independent public databases, yielding 5,714 curated pairs. For each of the 67 MSC-expressed ligands, we identified cognate receptors and systematically assessed their specificity to the B-cell lineage by cross-referencing expression across 11 non-B-cell BM subpopulations. Broadly expressed receptor-ligand pairs were excluded to maximize lineage specificity (Supporting Information S1: Figure 3). Candidate interactions were further prioritized by scoring the enrichment of each ligand’s predicted downstream target gene module at the single-cell level, retaining only those pairs where the cognate receptor was co-expressed in cells exhibiting positive module enrichment. Putative regulators were then cross-referenced against Gene Ontology terms encompassing B-cell and hematopoietic differentiation to identify biologically established factors within the candidate list. Database sources and analytical parameters are detailed in the Supplemental Methods.

**Figure 3:**
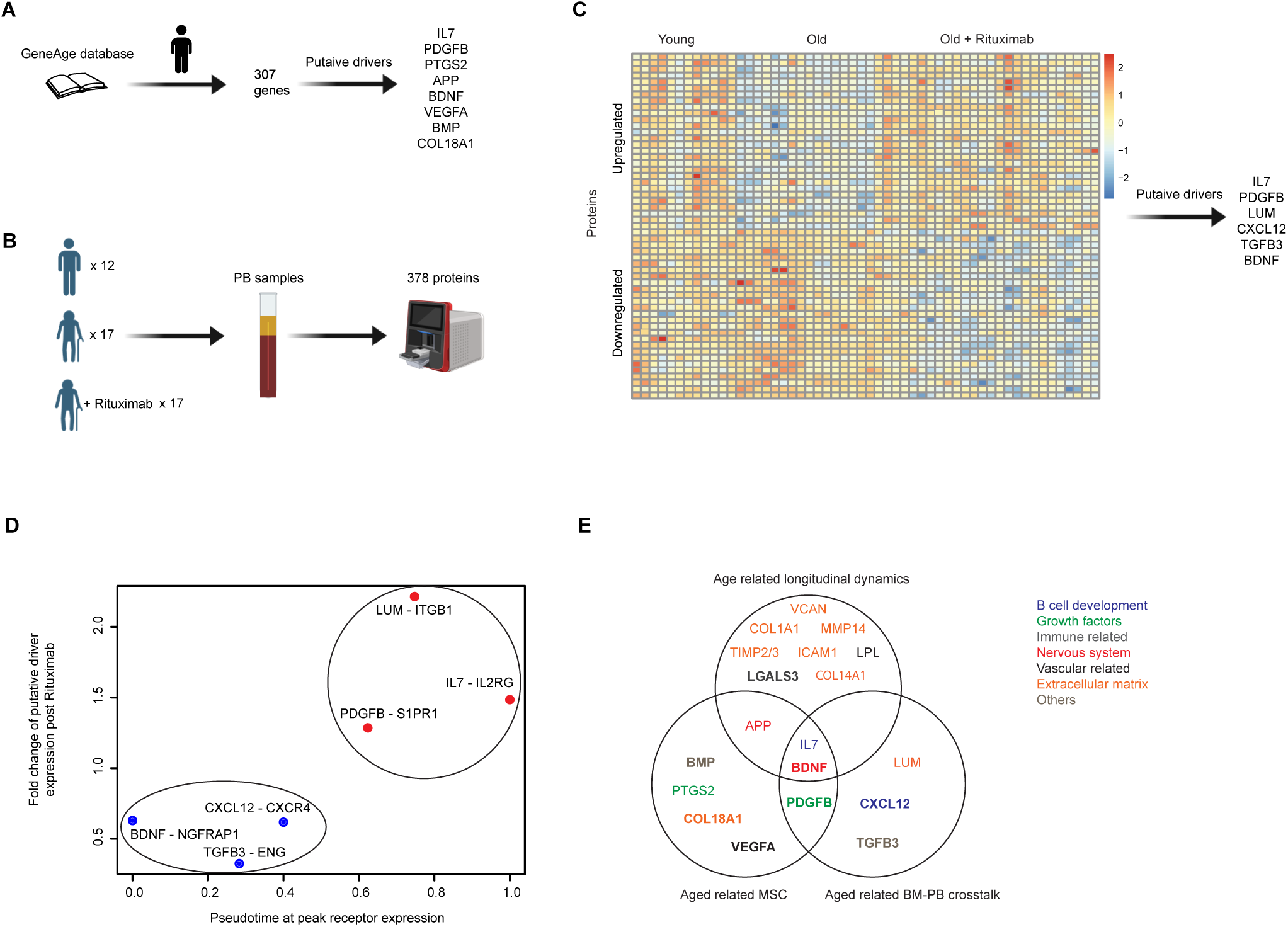
Prioritized candidate factors participate in age related regulation of B-cell development. **(A)** An illustration depicting a strategy to identify putative regulators with age-related expression in older individuals utilizing the GeneAge dataset. **(B)** An experimental design for exploring peripheral blood (PB) proteins during induction of B lymphopoiesis, whereby a multiplex protein assay was utilized to quantify the levels of 378 proteins in the peripheral blood plasma of young individuals (n=12), old individuals (n=17) and old patients that were treated with B-cell depleted therapy, Rituximab (n=26) **(C)** A heatmap showing the expression of soluble proteins whose profile in older patients treated with Rituximab was significantly altered(p.adjust < 0.05). Low and high normalized, scaled expressions are denoted as blue and red respectively. **(D)** A scatter plot demonstrating the correlation between B-cell developmental receptor activity along the trajectory (x-axis) and the behaviour of their paired, cognate ligands during induction of B lymphopoiesis (y-axis). **(E)** A Venn diagram of age-associated, prioritized putative regulators according to three aged related criteria: (i) shifted expression dynamics; (ii) regulation by mesenchymal stromal cells; and (iii) regulation by PB-BM cross talk. Candidates are colored differently in accordance to their known biological functions.

#### Mapping differences in longitudinal B lymphopoiesis during aging

To determine which candidate regulators were functionally relevant to age-related B-lymphopoiesis dynamics, we analyzed an independent published aging cohort (GSE120446; n=16 donors; 62,178 cells). Following normalization, dimensionality reduction, and cell type annotation, four progenitor B-cell subpopulations were isolated. Cell numbers were equalized across samples to mitigate inter-sample bias, and a consensus pseudotime trajectory was generated. Age-stratified trajectories were compared algorithmically to identify regulatory candidates whose expression patterns diverged with aging. Donor demographics, annotation validation, and pipeline parameters are provided in the Supplemental Methods and Supporting Information S2: Table 1.

### Biological Methods

#### Study cohorts and ethical approvals

Soluble protein profiling was performed on plasma collected from three clinically defined cohorts: healthy young adults (ages 18–35), healthy older adults (ages ≥55), and older patients (ages ≥55) with B-cell non-Hodgkin lymphoma previously treated with Rituximab. The study was approved by the Institutional Review Board (IRB 3097); inclusion and exclusion criteria have been previously reported ^55^. Fresh BM samples for functional experiments were obtained from healthy allogeneic BM donors at Rambam Medical Center (IRB 0086-21).

#### Soluble protein profiling

Differential plasma protein expression across the three cohorts was quantified using a Luminex bead-based multiplex platform. Group differences were assessed by parametric or non-parametric testing as appropriate following normality assessment, with correction for multiple comparisons. Significance was set at a corrected p-value threshold of p.adjust < 0.05. Full assay and statistical details are provided in the Supplemental Methods.

#### Functional validation: Single-cell gene expression stimulation assay

To functionally validate the candidate regulators identified computationally, we conducted a stimulation experiment in primary human BM cells. CD34+ BMMCs were isolated from healthy donors, combined with CD34-BMMCs, and cultured for 24 hours in the presence of B-cell developmental cytokines alongside candidate regulatory factors (detailed in Supporting Information S2: Table 3). Cells were then profiled by scRNA-seq. Post-stimulation transcriptomic data were normalized, dimensionally reduced, and annotated against a BM reference. Differentially expressed genes were identified using a scRNA-seq-appropriate framework with conservative multiple testing correction, and pathway enrichment was assessed against the Reactome database. To quantify transcriptional similarity across stimulation conditions, Jaccard similarity coefficients were computed for enriched pathway pairs across cell subpopulations. Cellular proliferative and interferon response activity was additionally quantified per sample using single-sample gene set enrichment analysis (ssGSEA) against curated gene sets. Full library preparation, sequencing, demultiplexing, and bioinformatic pipeline details are provided in the Supplemental Methods and Supporting Information S2: Table 2.

#### Functional validation: *In Vitro* B-cell development assay

To specifically validate the role of BDNF in supporting B-cell development, we employed an established stromal co-culture system in which CD34+ BMMCs were grown over a confluent HS-5 stromal cell layer for three weeks. Cells were harvested at defined time points and immunophenotyped by flow cytometry. Gating strategies, staining panels, and culture conditions are described in the Supplemental Methods and Supporting Information S1: Figure 4 and Supporting Information S2: Table 4.

**Figure 4:**
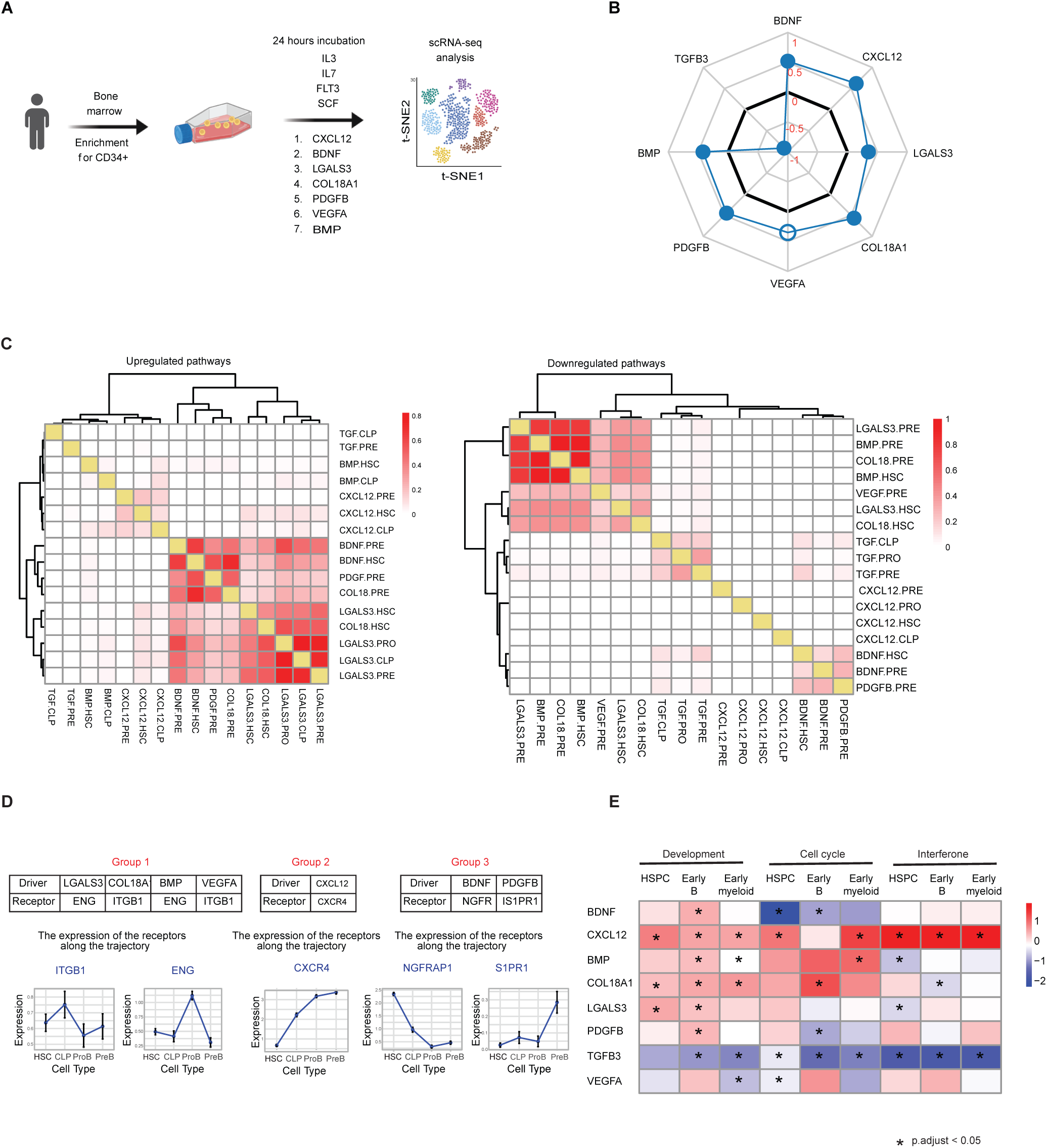
Single cell stimulation assays suggest distinct mechanisms underlying the possible effects of the putative regulators on B lymphopoiesis. **(A)** A schematic depicting the experimental design for testing stimulation effects of putative B-cell developmental regulators. Healthy, human CD34^+^ enriched cells were incubated for 24 hours in the presence of both cytokines, required for survival and differentiation of B cells, and each of the indicated putative regulators, followed by scRNA-seq. **(B)** Candidates differentially affect biological pathways in progenitor B cells. A radar chart showing the enrichment score of a gene set comprising 122 genes associated with cell growth, cell proliferation and transcriptional regulation (Methods) following simulation with the putative regulators. The thick black line represents a neutral effect on the pathway (0). Filled and empty circles represent statistically significant and non-significant results, respectively. **(C)** A heatmap showing Jaccard similarity levels between significant pathways upregulated (left) and downregulated (right) relative to control samples, following stimulation with the annotated putative regulators. **(D)** Summary of three regulator groups clustered by their post-stimulation behaviors, as assessed by Jaccard similarity analysis. Each group demonstrates distinct dynamics in their paired receptors along the trajectory. **(E)** A heatmap demonstrating the enrichment score of developmental, cell cycle and interferon related pathways within HSPCs, progenitor B and myeloid subpopulations, as obtained from ssGSEA analysis.

## Results

### Mapping intercellular communication in human bone marrow identifies candidate factors predicted to influence B-cell development

In order to identify regulators of B lymphopoiesis, we took a three-pronged approach (**Figure 1A**): first, to initially generate a list of candidate factors, we explored intercellular communication networks between progenitor B cells and MSCs in the BM, since MSCs are master regulators of B-cell differentiation ^20,56^ ; second, we prioritized these subsequent, candidate factors by leveraging differences present between young and aged B lymphopoiesis, as aging is a model in which B-cell development is naturally diminished; and third, we validated our putative regulators using scRNA-seq and an *in-vitro* system of B-cell development.

Initially, we sought to map nascent progenitor B-cell genes to identify genes whose dynamic interplay orchestrates the development of B cells. To that end, we visualized the cells by UMAP, using publicly available data (GSE149938) ^57^ and revealed distinct cell populations arising from HSPCs in line with the known hierarchy of BM maturation (**Figure 1B**). We further constructed a high-resolution scRNA-seq trajectory, recapturing the known order of B-cell differentiation, with cells positioned along a continuum of pseudotime as opposed to discrete stages, reflecting the underlying biology more accurately (**Figure 1C**). To validate the trajectory, we characterized the expression dynamics of genes whose expression levels vary throughout the development of B cells and confirmed that their expression profiles along the trajectory are as expected and are supported by the literature ^52^ (**Figure 1D**). Of note, we removed from consideration any genes whose expression along the trajectory was observable yet uniform (i.e. not distinct to a single B-cell subpopulation), which resulted in 1,381 genes with dynamical expression patterns and potential contributions to the B-cell developmental trajectory, from hereon referred to as pseudotime associated genes (Supporting information S2: Table 5).

To characterize the microenvironment of developing B cells - a crucial aspect of differentiation control - we mapped the transcriptional profile of MSCs (Supporting information S2: Table 6). Subsequently, to explore interactions between developing B cells and MSCs, we integrated three separate databases of receptor-ligand pairs, curating a comprehensive list of human receptor-ligand interactions ^58–60^. Importantly, this list included pairs that mediate interactions between different cells, as inferred from the literature or as shown previously in biological experiments (Supporting information S2: Table S7). We utilized the receptor-ligand list in-conjunction with the scRNA-seq datasets to search for putative regulators, defined as ligands that are expressed by MSCs and interact with receptors that are expressed by B-cell progenitors, according to the following stepwise process (**Figure 1E**): (i) out of the 1,381 B-cell pseudotime associated genes, we identified 101 receptors dynamically expressed along the developmental trajectory; (ii) we identified 389 ligands that potentially interact with those receptors based on our assembled interaction list; (iii) of those 389 ligands, 67 ligands are expressed by MSCs in the BM; (iv) of these, we identified 58 MSC ligands with high specificity to progenitor B cells; and finally (v) we included a downstream analysis step and identified 52 ligands whose downstream target genes and cognate receptors are expressed by the same progenitor B cells. Collectively, these ligands made up our initial list of candidate factors predicated to act on B lymphopoiesis for further investigation. By utilizing GO annotations, we identified only 5 genes associated with known human hematopoietic and B-cell developmental/differentiation pathways (Supporting information S2: Table 8). Thus, the remaining 47 candidate factors are seemingly related to different and unrelated biological functions (**Figure 1F**). This lack of clarity led us to further search for ways to test their possible role in the B lymphopoiesis process and narrow down the list of candidate factors for testing.

### Pairwise trajectory alignment reveals age-dependent differences in the expression dynamics of pseudotime associated genes

To further prioritize candidate factors and identify putative regulators of B lymphopoiesis, we searched for conditions in which the developmental process is naturally perturbed, reasoning that factors associated with such deviations are more likely to play a functional role in proper development. As such, we focused on aging, the most common and accessible condition. Since it is unlikely that key genes are completely absent in the elderly, who still produce low levels of B cells, perturbations of other kinds may instead be responsible for loss of B cells with age. Therefore, we decided to explore the differences in the longitudinal development of aged B lymphocytes. To do so, we initially analyzed a publically available scRNA-seq dataset of 16 healthy BM donors aged 24 to 67 years old ^61^. To overcome baseline differences between the samples, we created a common B-cell trajectory that included all the donors, termed here as the “consensus trajectory”. We compared the dynamics of each of the previously identified pseudotime-associated genes across the different ages relative to the consensus trajectory by using the cellAlign trajectory-alignment algorithm ^62^ (**Figure 2A**). Per gene, this algorithm identifies differences in gene expression dynamics between two independently constructed single cell trajectories by quantifying the pseudotime shift between. To avoid technical obstacles resulting from baseline pseudotime shifts between young and old that may provide false positive age-associated dynamic genes, we performed 56 pairwise comparisons within the same age groups (28 comparisons within 8 young donors and 28 comparisons within 8 old donors) and 64 comparisons across age groups (**Figure 2B**). Subsequently, we identified only a few genes with significant pseudotime shifts across the comparisons (p.adjust < 0.05). In contrast, as age introduced gene expression heterogeneity ^63^, we omitted comparisons performed within older donors, which revealed 399 genes with significantly different expression dynamics in pairwise comparisons within the young samples vs. pairwise comparisons across age groups (p.adjust < 0.05) (**Figure 2C**). Repeating this on permuted groups resulted in the identification of 257 genes with significant pseudotime shifts (p.adjust < 0.05) (**Figure 2D,E**). To identify which candidate factors also display age-related shifts in expression dynamics, we intersected the results with our receptor-ligand and candidate factors lists and found that out of the 257 age-related genes, 5 pseudotime related receptors: CD74, NGFRAP1, CD44, SPN and IL2RG were present. Accordingly, we prioritized the cognate ligands of these receptors: APP (interacts with CD74), BDNF (interacts with NGFRAP1), COL14A1, COL1A1, VCAN, TIMP2/3, MMP14 and LPL (interact with CD44), ICM1 and LGALS3 (interact with SPN) and IL7 (interacts with IL2RG). In summary, this step resulted with 11 ligands that may be involved in B-cell development.

### Prioritized candidate factors participate in age related regulation of B-cell development

To further bolster our analysis, we next explored the differential, inter-cellular regulation of B lymphopoiesis upon aging. Initially, we utilized “GenAge”, the aging gene database, to extract aging-related differentially expressed genes in MSCs. This yielded 8 MSC ligands/putative B-cell regulators with differential expression upon aging within humans: PDGFB, VEGFA, APP, IL7, BDNF, PTGS2, Collagen 18A and BMP (**Figure 3A**).

Beyond studying the direct regulation of B lymphopoiesis by aged MSCs, we also explored the regulation of B lymphopoiesis orchestrated by cross-talk between mature peripheral blood B lymphocytes and progenitor B cells in the BM - a process known to be responsible for replenishing the B lymphocyte pool upon aging ^64,65^. Previous studies have already shown that the removal of circulating B cells by B-cell depletion therapy in elderly individuals stimulates their production in the BM ^55,66,67^. Therefore, to further identify and prioritize putative regulators that change during stimulation of B lymphopoiesis in elderly individuals, we tested the differential regulation of plasma proteins between young individuals, old individuals and old individuals treated with the B-cell depletion therapy, Rituximab, administered due to Non-Hodgkin Lymphoma (NHL) (**Figure 3B**). Following analysis of plasma soluble proteins using a 378 Luminex protein panel, we identified 55 plasma proteins whose expression in older patients, treated with Rituximab (i.e. during the B lymphopoiesis process), was altered relative to untreated old individuals, suggesting a role in B lymphopoiesis (p.adjust < 0.05) (**Figure 3C**). Out of these factors, we prioritized six soluble proteins: three putative regulators whose level increased following treatment with Rituximab (IL7, PDGFB and LUM) and three putative regulators whose levels decreased following treatment with Rituximab (CXCL12, TGFB3 and BDNF). Interestingly, we found that the putative regulators that were upregulated in older patients treated with Rituximab also interact with receptors whose expression is higher at later stages along the trajectory. In contrast, downregulated putative s interact with receptors expressed at the beginning of the trajectory (**Figure 3D**). Taken together, by analyzing the regulation of B lymphopoiesis in older individuals, we prioritized ∼40% of the putative regulators that participated in age related interactions and which displayed diverse biological functions (**Figure 3E**). Importantly, IL7 and BDNF were identified by all three aspects of this screening process. Notably, and consistent with our analysis, it has been shown that in humans IL7 is crucial for commitment to the B-cell lineage and for proliferation of early human B-cell progenitors ^32^, demonstrating the robustness of our process.

### Single cell stimulation assays suggest distinct mechanisms underlying the possible effects of each putative regulator on B lymphopoiesis

To obtain evidence regarding the possible, functional role(s) of each putative regulator, we performed a single cell gene expression stimulation assay. Specifically, we sought to investigate whether putative regulators affect B-cell development *in-vitro,* as suggested by our previous analyses, and if so, to determine which subpopulations are affected and highlight the underlying mechanisms. For this, we incubated healthy human BM cells enriched for CD34^+^ cells supplemented with B-cell maintenance cytokines, along with each of the putative regulators separately for 24 hours prior to single cell sequencing (**Figure 4A**). We prioritized putative regulators where: (i) their role in human B lymphopoiesis has not yet been established; (ii) they were prioritized by at least one of the age related criteria listed above; and (iii) represented diverse biological functions, ultimately selecting: LGALS3, COL18A1, BMP, TGFB3, BDNF, VEGFA and PDGFB. CXCL12 was utilized as a positive control. To assess the possible functional role of putative regulators in the context of B-cell development, we compared gene expression levels and pathway activity scores following each putative regulator stimulation to the control sample within the HSPC and progenitor B cellular compartments. Consistent with expectations and prior literature, CXCL12 upregulated immune related pathways within progenitor B-cell (Supporting information S1: Figure 5A). To gain insight into the possible role each,different stimulation may have on B-cell development, we tested their ability to upregulate development-related genes - that is, a group of genes previously associated with cell growth, proliferation and transcriptional regulation and observed that BDNF, CXCL12, LGALS3, COL18A1, PDGFB and BMP significantly upregulated these pathways on early B cells (p.adjust < 0.05), suggesting they may play a role in B lymphopoiesis (**Figure 4B**). By contrast, TGFB3 down-regulated immune and developmental-pathways, and therefore was eliminated from further consideration as a lymphopoietic regulator (Supporting information S1: Figure 5B). VEGFA showed elevated trends in developmental related genes, but failed to reach statistical significance. To identify commonalities between stimuli, and obtain a comprehensive view of the effects we computed Jaccard similarity coefficients between pairs of biological pathways within subpopulations that were significantly changed relative to the control for each stimulation (**Figure 4C**). While taking into consideration both up and down-regulated biological functions, we identified 3 groups that exhibit shared, similar behaviour post stimulation (Group 1: LGALS3, COL18A1, BMP and VEGFA. Group 2: CXCL12. Group 3: BDNF and PDGFB). Interestingly, each group also possesses distinct receptor expression dynamics along the B-cell developmental trajectory. The receptors of LGALS3, COL18A1, BMP and VEGFA peak at the CLP-ProB phases, in the middle of the trajectory (**Figure 4D**). C-X-C Motif Chemokine Receptor 4 (CXCR4), the receptor of CXCL12, is detected at low levels in HSPCs and its expression gradually increases toward the terminal end of the process. Although the receptors of the third group harbored different and inconsistent patterns relative to those observed in the first two clusters, they demonstrate opposing dynamics to one another: while the expression of NGFRAP1, that binds to BDNF, is higher at the beginning of the trajectory, the expression of S1PR1, PDGFB’s receptor, peaks toward the end of the trajectory despite the similarity in gene expression changes induced by BDNF and PDGFB.

**Figure 5:**
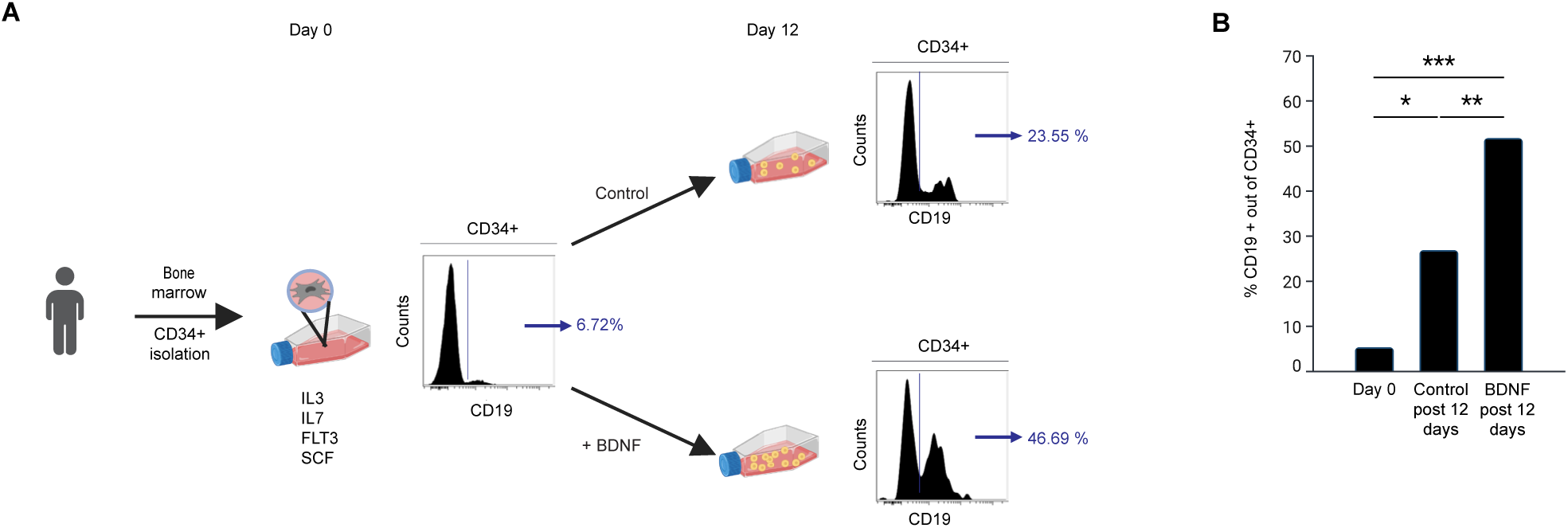
BDNF enhances the differentiation of B cells in a co-cultured biological system. **(A)** A schematic depicting the experimental design aiming to test the impact of BDNF on HSPCs in an *in-vitro* biological system. CD34^+^ cells were isolated from healthy human bone marrow samples and co-cultured on SF-5 cells layers with the B-cell related cytokines, IL3, IL7, FLT3 and SCF, as well as BDNF. After 12 days, cells were harvested and analyzed by flow cytometry. Representative plots of CD19^+^ cells out of CD34^+^ cells at baseline and following 12 days of co-culture with and without BDNF are shown. **(B)** Bar plot showing the % of CD19^+^ cells out of CD34^+^ cells 12 days post co-culturing with BDNF.

Given the activation of similar biological pathways as revealed by Jaccard similarity analysis, we sought to further assess the exact nature of the biological functions influenced by the different stimulations, and inspected the pathway alterations following the different stimulations **(Figure 4E**). We found that in contrast to CXCL12 and BMP stimulations, both BDNF and PDGFB significantly downregulated the cell cycle pathway. While upregulation of the cell cycle pathway is crucial for proliferation, a quiescence state is required for further differentiation. Specifically, during the development of early B cells, quiescence is necessary for the variable-diversity-joining (VDJ) recombination process ^52,68^. VDJ recombination randomly rearranges gene segments to create a diverse repertoire of heavy chain variable regions, thus enabling the immune system to recognize and respond to a wide range of pathogens, whereas at later stages, signals from the pre-B cell receptor (BCR) re-establish quiescence to facilitate recombination of the immunoglobulin light chain ^69,70^. In line with that, both BDNF and PDGFB significantly upregulated the RNA binding protein, ZFP36L2, which was shown to have a critical role in promoting VDJ and immunoglobulin light chain recombination by maintaining quiescence ^71^. Thus, our data suggests that both BDNF and PDGFB promote B lymphopoiesis by arresting the cell cycle and promoting quiescence, however, their different cognate receptor expression dynamics (NGFRAP1-BDNF, high at the beginning of the trajectory, and S1PR1-PDGFB peaking post pre-BCR) suggested they may act at different developmental stages to accomplish a similar task. Interestingly, while the interferon pathway was upregulated following stimulation with CXCL12, it was downregulated post stimulation with LGALS3, COL18A1, VEGFA and BMP. Interferons (IFNs) were initially characterized based on their crucial role in the innate antiviral immune response ^72^. However, subsequently their role in inhibition of cell growth ^73–75^ and in balancing the immune response ^76,77^ has been established. Therefore changes in interferon signaling following induction of B lymphopoiesis may be expected and point to the different mechanisms that are mediated by IFNs during B-cell development following the stimulations. Upregulation of IFN related pathways by CXCL12 points to its role in promoting immune function. However IFN downregulation following LGALS3, COL18A1, VEGFA and BMP stimulation, suggests maintaining the balance of immunity and the prevention of autoimmunity, may also be important during B-cell development. Finally, we observed that BDNF, LGALS3 and PDGFB stimulations were specific to the B-cell lineage, as they did not activate pathways in myeloid cells. In summary, by testing the effects of the different stimulations from a functional activation viewpoint and along the course of B-cell development, our analysis suggests different mechanisms by which they may regulate the process of B lymphopoiesis.

### BDNF enhances the differentiation of B cells in a co-culturing biological system

While our single cell stimulation assays revealed that the tested B-cell regulators may promote B-lymphopoiesis in a myriad of ways, B-cell development is ultimately a longer process than the 24-hour window we accommodated for in our experimental design, and any regulator must yield an increase in proliferating B cells to truly demonstrate regulation capacity and therapeutic potential. To advance our hypotheses further, we focused on a single putative regulator and an orthogonal experimental design, an *in-vitro* cell system, to further test the plausibility that we have identified a B-cell developmental regulator. To that end, we chose BDNF for further validation, due to: i) it being the only putative regulator (except for IL7, whose role in B lymphopoiesis has already been established ^34^) to register positive across the three age-related analyses (**Figure 3E**); ii) previous studies showing the possible role of BDNF in mice ^78,79^, suggesting that BDNF has the ability to affect the B lymphopoiesis process in humans as well; and (iii) BDNF acting at the beginning of the trajectory (Supporting information S1: Figure 6), making it an attractive and appropriate therapeutic agent following anti CD19/ anti CD20 therapy which eliminates early progenitor B cells.

We profiled BDNF effects in an *in-vitro* biological system that mimics the development of B cells, starting with HSPCs ^80^. For this, we isolated CD34^+^ human bone marrow mononuclear cells (BMMC) from healthy human BM samples and co-cultured them with HS5 stromal-cell feeders in the presence of IL3, IL7, SCF and FMS-like Tyrosine Kinase 3 (FLT3). Using flow cytometry analysis we detected a gradual increase in the number of HSPCs differentiating into progenitor B cells that express increasing levels of CD10 and CD19 over the course of 3 weeks (Supporting information S1: Figure 7, 8). As the expression of the BDNF receptor, NGFRAP1, is highest during the early developmental stages along the trajectory after which it gradually decreases, we hypothesized that the developmental effect of BDNF will be most pronounced in the early stages of the system. The appearance of CD19 expression on CD34^+^ cells marks their commitment to the B-cell lineage ^81–83^. Examination of the differentiation of CD34^+^ HSPCs at 12 days following addition of BDNF revealed that a larger proportion of the CD34^+^ HSPCs had differentiated into Progenitor B cells expressing CD19 (P = 0.0042) (**Figure 5 A,B**). Taken together, our multi-step prioritization approach - that took into consideration age related criteria and functional effects - led us to identify and successfully validate BDNF as a factor that can regulate or enhance B-cell development, in an *in-vitro* biological system.

## Discussion

Here, to address the unmet clinically important challenge of B-cell aplasia we leveraged an integrative, systems-level approach to identify and prioritize putative regulators for B lymphopoiesis. During recent years, in the post-genome-AI era and the explosion of molecular biology techniques, novel opportunities for drug discovery have emerged. ^84–87^. Many targets fail in pre-clinical and clinical trials due to a lack of efficiency and/or side effects. To optimize an early step of the drug discovery process, we chose candidate factors that were inferred from across several independent data sources,leveraged differences in an impaired version of the process under study (i.e. B-lymphopoiesis during aging), and utilized clinical samples to narrow down candidate factors. Ultimately, we suggest and hypothesize that by integrating different aspects and points of view on B lymphopoiesis, early during the process of drug discovery, we build confidence and elevate the probability of target success and/or successful translation into the clinic. Additionally, by testing the cell specificity of potential regulators early stages, our prioritization approach reduces the chance of side effects that may result from targeting unintended but crucial functions.

Our strategy highlighted a role for BDNF, the most prevalent growth factor in the central nervous system ^88–90^, in B-cell regulation. The concept of “from neurotrophins to neurokines” has been previously proposed, stating that neurotrophins, primarily recognized for their crucial involvement in neuronal development and survival, are also involved in immune system processes ^91^. Nevertheless, the role of BDNF in human immunity and specifically B-cell development remains largely unexplored. We show here for the first time that BDNF regulates the development of human BM B cells. However, and importantly, we are aware of the potential autoimmune diseases and immune dysregulations that may result from immune system-specific interventions and therefore the safety of agents, such as BDNF, should be thoroughly evaluated. Nevertheless, BDNF is currently utilized in clinical trials for the treatment of Alzheimer’s disease (NCT05040217), suggesting a low toxicity profile.

Exploring the influence of putative regulators throughout B-cell development yields interesting insights. Our observations indicated that in older patients treated with Rituximab, upregulated, putative regulators interacted with receptors expressed at later stages of the trajectory compared to those downregulated. Such an observation may be explained by the following mechanism: upon treatment with Rituximab, an anti-CD20 antibody, cells that do not yet express CD20 proliferate, leading to an increase in available receptors and a decrease in free ligands. Such a mechanism is consistent with our results: we observed decreases in ligands that interact with the early B-cell trajectory receptors, CXCL12, BDNF and PDFGB. In contrast, CD20 expressing cells are eliminated by the treatment, resulting in an increased presence of free ligands that interact with the late B-cell trajectory receptors: IL7, PDFFB and LUM (Supporting information S1: Figure 8).

Progress towards acceleration of human B-cell lymphopoiesis in patients will advance in a stepwise manner and there are multiple additional steps to be taken before a ‘first-in-human’ trial for B-cell lymphopoiesis reconstitution with BDNF or any of the other putative regulators discovered, can occur. Further evidence can be generated through additional single cell and functional studies, the testing of additional regulators, in combinations, and in more diverse conditions. Of particular note, it is important to question the utility of validating such complex system-level regulators in animal models, where much of the underlying disease biology is not fully conserved ^48–50^. In summary, through our comprehensive approach that integrated different types of data - from detailed genetic analysis, to blood samples and biological experiments - we looked at the process of B lymphopoiesis from different angles simultaneously. Such an approach may provide a framework for studying complex developmental processes in the BM, and in other tissues, and pave the way to discover accelerators for other immune compartments as well toward improving immune system function.

## Supporting information

Supportive information S1

Supportive information S2

Graphical abstract

## Acknowledgments

Shai S. Shen-Orr acknowledges gracious support for this work by NIH NIAID award PO1 A153559-01.

Neta Nevo was supported by the Rubenstein Technion Integrated Cancer Center Fellowship.

