## Supplementary material for "A systems approach to B-cell development identifies BDNF as a regulator of human B lymphopoiesis": Supportive information S1

**Additional file 1: Supplemental Methods**

**Additional file 2: Supplemental Figures**

#### Additional file 1: Supplemental Methods

##### scRNA-seq preprocessing and quality control

Published scRNA-seq data (GSE149938) were imported into R [v4.3.1]. Preprocessing and quality control were performed as previously specified [1]. Cell type annotations were validated by plotting the expression of canonical B-lineage markers, CD34, MME (CD10), CD19, MS4A1 (CD20), CD79A, and TNFRSF17 (BCMA), across clusters using Seurat's ViolinPlot() function [v4.0.3] [2], confirming a biologically consistent progressive expression pattern from HSPCs to plasma cells (Supporting Information S1: Figure 1a). A total of 18,008 genes across 7,643 cells were analyzed.

Normalization and scaling were performed using SCTransform via Seurat [v4.0.3]. Dimensionality reduction used the RunPCA and RunUMAP functions in Seurat under default parameters, retaining the top 3,000 variable features and 25 principal components to generate the UMAP embedding. Visualizations were produced using dynplot [v1.1.1] [3].

#### Differential gene expression analysis

Differential expressed genes were identified using the MAST algorithm [v1.18.0] via Seurat's FindMarkers function. Adjusted p-values were calculated using the Benjamini-Hochberg (BH) correction procedure; genes with p.adjust < 0.05 were retained for downstream analysis.

#### Trajectory inference and gene dynamics modeling

Trajectories were inferred using SCORPIUS [v1.0.9] [4] under default parameters. Gene expression dynamics along trajectories were modeled using tradeSeq [v1.20.0] [5] via the fitGAM and associationTest functions, which assess non-parametric association between averaged gene expression and pseudotime without assuming a fixed dynamical shape. Two key outputs were used: (1) the Wald statistic, assessing whether pseudotime-associated model coefficients differ significantly from zero; and (2) the mean log2 fold change (MeanLogFC) across the trajectory. Genes with a Wald statistic > 120 were first selected (n = 1,556). This threshold was chosen from the inflection point of the Wald statistic distribution curve, where gene counts dropped sharply, marking the transition from background noise to biological signal (Supporting Information S1: Figure 2). Combined with the BH-corrected DEG filter (p.adjust < 0.05), these steps retained 1,381 genes from an initial 3,914. A deliberately lenient threshold was applied at this stage, as subsequent biologically informed filtration steps were planned.

#### Receptor-ligand database integration

Three independent receptor-ligand databases were integrated:

1. **NicheNet (v2):** Lr_network_human_21122021.rds - 4,986 pairs [6]
2. **GCNG:** 41467_2015_BFncomms8866_MOESM611_ESM (Version 1) - 2,557 pairs [7]
3. **Qiao et al.:** Msb145141-sup-0011-table2 - 933 pairs [8]

Duplicate pairs appearing in more than one database were removed and sources unified. The final list comprised 5,714 unique pairs, including 1,298 unique ligands and 1,236 unique receptors.

**Receptor-ligand lineage specificity analysis**

For each of the 67 MSC-expressed ligands, cognate receptors were identified using the integrated list above. Top marker genes for each of the 11 non-B-cell BM subpopulations were identified using FindMarkers(); genes with p.adjust < 0.01 and average log2FC > 1 (i.e., specific to non-B-cell lineages) were excluded. Receptor expression levels for each ligand were then cross-referenced across all non-B-cell subpopulations, and nine ligands whose defined receptors were expressed in more than six non-B-cell subpopulations were excluded **(Supporting Information S1: Figure 3)**.

**Downstream target gene scoring and interaction prioritization**

To infer the probability that specific receptor-ligand interactions have occurred, we utilized NicheNet's downstream target gene matrix (https://zenodo.org/record/3260758/files/ligand_target_matrix.rds) and cross-referenced this with receptor expression patterns in progenitor B-cells. Specifically, for each ligand, we construct a module of the 20 highest-probability downstream target genes. This range was tested from 10 to 40 downstream genes with consistent results. Module enrichment was scored at the single-cell level using Seurat's AddModuleScore function. Ligands where >50% of cells within a given subpopulation exhibited a positive module score were identified, and receptor-ligand pairs were retained only when the cognate receptor was co-expressed in that same subpopulation.

**Gene ontology annotation of putative regulators**

Putative regulator genes were cross-referenced against the following GO terms, selected in a supervised manner based on prior biological knowledge:

| GO Term | Description |
|---|---|
| GO:0002327 | Immature B-cell differentiation |
| GO:0071425 | Hematopoietic stem cell proliferation |
| GO:0001923 | B-1 B-cell differentiation |
| GO:0001924 | Regulation of B-1 B-cell differentiation |
| GO:0045577 | Regulation of B-cell differentiation |
| GO:0030183 | B-cell differentiation |
| GO:0045579 | Positive regulation of B-cell differentiation |
| GO:0001926 | Positive regulation of B-1 B-cell differentiation |

| | |
|---|---|
| GO:0001925 | Negative regulation of B-1 B-cell differentiation |
| GO:0060218 | Hematopoietic stem cell differentiation |

**Age-associated B-lymphopoiesis: data processing**

Published data from GSE120446 were imported into R [v4.3.1]. Donor codes and demographic characteristics for all 16 donors are provided in Supporting Information S2: Table 1. In total, 13,112 genes across 62,178 cells were analyzed. Normalization and scaling used SCTransform via Seurat [v4.0.3]. Dimensionality reduction used RunPCA and RunUMAP with the top 500 variable features and 10 principal components.

Cell type annotation used the SingleR function [9] with two reference datasets: HumanPrimaryCellAtlasData and OvershternHematopoieticDataN. Annotations were validated using VlnPlot() in Seurat, confirming progressive expression of CD34, MME (CD10), CD19, MS4A1 (CD20), CD79A, and TNFRSF17 (BCMA) consistent with biological expectations (Supporting Information S1: Figure 1B). Four progenitor B-cell subpopulations were isolated; cell numbers were equalized across samples to control for inter-sample bias. Samples were merged and a consensus pseudotime trajectory was generated using SCORPIUS. Age-stratified trajectories were compared against the consensus trajectory using the CellAlign algorithm [10] .

**Soluble protein profiling: Luminex multiplex assay**

Data were collected on a Luminex 200 instrument and analyzed using Analyst 5.1 software (Millipore). Median Fluorescence Intensity (MFI) values were used as the primary readout. Samples with a mean bead count of <50 were excluded. MFI values were calculated by blank reduction.

Between-group differences were tested using one-way ANOVA with Tukey's multiple comparisons correction when sample distributions were normal per Shapiro-Wilk testing, or by Kruskal-Wallis test with Dunn's multiple comparisons correction when distributions were non-normal. All results were adjusted for multiple comparisons using the BH procedure, with p.adjust $< 0.05$ considered statistically significant.

**Single-cell gene expression stimulation assay**

RNA single-cell library preparation and sequencing was conducted by the Technion Genomics Center, Technion - Israel Institute of Technology, Haifa, Israel.

Sample Fixation and Processing

Samples were fixed using the ScaleBio Sample Fixation Kit (ScaleBio, cat# 2020001) according to the manufacturer's protocol. Fixed cells were stored at -80°C overnight and thawed the following day. Subsequently, the cells were processed according to the ScaleBio post-storage protocol. Cells were diluted to a working concentration of approximately 2,000 cells/µl.

Library Prep

Fixed RNA single-cell libraries were prepared using the ScaleBio single-cell RNA Sequencing Kit v1.1 (Scalebio, cat# 950884) according to the manufacturer's protocol. 5 ul of diluted cell suspension (a total of 10,000 cells) was distributed into each well of the RT Barcode Plate. After the ligation step and before final distribution, all cells were counted again and diluted according to the protocol, yielding a final concentration of 355 cells/ul. Final library QC was performed by measuring library concentrations using Qubit (Invitrogen) with the Equalbit dsDNA HS Assay Kit (Vazyme, cat no. EQ121) and size was determined using the TapeStation 4200 (Agilent) with the High Sensitivity D1000 kit (Agilent, cat no. 5067-5584). The library exhibited a peak with an average size of approximately 377 bp, indicating a robust, fixed RNA profiling library trace.

Sequencing Details

Sequencing was performed on the Illumina NextSeq 2000 platform, using P4 XLEAP-SBS Reagent Kit 100 cycles (Read1-100; Index1-8; Index2-8 ; Read2-0) (Illumina, cat no. 20100994).

Demultiplexing, Mapping and Cell Calling

Demultiplexing of the sequencing run was performed for all scRNA-Seq libraries using bcl2fastq (v2.20.0.422), utilizing dual indices with 1 mismatch allowed. The barcodes used for demultiplexing were according to the RNA-A-AP1 plate found in the samplesheet_v1.1_revComp file - as downloaded from the ScaleRna pipeline github page. In the sequencing run, 1,698,847,591 reads passed the filter. Out of which, 1,311,034,079 reads (77.2%) were allocated to any of the RNA-A-AP1 indices. Demultiplexed reads, organized in 96 sets of R1, R2 & I1 FASTQ files, were used as input for the ScaleRna pipeline. Following demultiplexing, primary analysis was performed by running the ScaleRna pipeline (v1.5.0) with the default parameters in the "conda" configuration, using NextFlow (v23.04.3).

Out of the sequencing run, 838 million reads were associated with one of the 12 samples. The distribution of reads among samples ranges between 47M to 90M reads in total. Around 93% of reads were mapped to the reference genome in every sample and ~84% of reads were mapped to the transcriptome. The pipeline yielded 167,363 cells overall, distributed between samples ranging from 9,957 to 17,857 cells (5.9% - 10.7%, expected = 8.3%) per sample. Percent of reads in cells averaged around 90% in all samples. The median unique transcript count per cell ranged, per-sample, between 1,236 to 1,459 with a median genes detected range between 1,003 to 1,161. These metrics are associated with library size while considering percent of reads in cells (Supporting information S2: Table 2).

**Post-Stimulation scRNA-seq analysis**

Normalization and scaling used SCTransform via Seurat (v4.0.3). Dimensionality reduction used RunPCA and RunUMAP with the top 500 variable features and 10 principal components. Cell types were annotated using the BM reference ("bonemarrowref") via Seurat's Azimuth runAzimuth function.

DEGs were identified using MAST (v1.18.0) via Seurat's FindMarkers with Bonferroni-adjusted FDR correction; a gene was considered differentially expressed at $|\log2FC| > 0.30$. Pathway over-representation analysis used clusterProfiler (v4.0.0) against the Reactome database, retaining pathways at FDR < 0.01 (Bonferroni correction). Jaccard similarity coefficients were computed for all pairs of enriched pathways per stimulation condition and cell subpopulation, defined as the ratio of pathway gene set intersection to union.

ssGSEA was performed using the GSVA R package (v1.50.5) against three curated gene sets:

1. Proliferation: Curated from Reactome and MSigDB pathways covering DNA replication, mitotic progression, chromosomal segregation, and canonical proliferation regulators: CDC45, CLASP2, MIS12, CEP250, ANKLE2, TOP3A, PCNT, NSD2, UBE2V2, PRIM1, RAD21, PPP2R2D, PRKAR2B, NUP93, CDCA5, DYRK1A, LBR, CCNA2, SMC2, MCM8, CEP41, MCM10, SKA1, CENPK, SKA2, KIF18A, LIG1, ODF2, MNAT1, PKMYT1, CSNK2B, NUP155, RNF168, MIS18A, ANAPC1, MCM7, TMPO, NUP107, HAUS5, E2F6, CEP192, MLH1, BRCA1, CHEK1, CDT1, KNTC1, POLR2E, FEN1, BLM, CCP110, GMNN, SYNE2, GINS1, CUL1, SMC4, TERF2, CKAP5, AURKB, KIF23, SGO2, CDC20, PSMB5, CDK5RAP2, RFC1, HAUS7, CEP152, PSMD14, KPNB1, CENPX, NDC80, CENPU, NCAPD3, CDC25B, TFDP1, TYMS, XPO1, ORC3, RBL1, BUB1B, RRM2, CDC25A, RANBP2, CLSPN, RAD51, NCAPG, YWHAH, MYBL2, PSME3, HJURP, BARD1, CEP78, DHFR, POLA1, CCNE1, SMC1A, POLD3, POLE2, DNA2, BRCA2, MND1, HMMR, TPX2, TERT, CDC7, CDCA8, VRK1, NUP153, BRIP1, BUB1, NCAPH, RFC3, KNL1, PRIM2, NCAPG2, CENPE, EXO1, TUBA1B, NSL1, TUBB, CENPF, MCM4, LMNB1, TOP2A.
2. Cell cycle**:** REACTOME_CELL_CYCLE_MITOTIC
3. Interferon signaling**:** REACTOME_INTERFERON_SIGNALING

**Biological Methods**

**BM sample acquisition and processing**

Fresh healthy human BM samples were obtained from allogeneic BM donors at Rambam Medical Center (IRB 0086-21). BMMCs were isolated by Ficoll density gradient centrifugation (STEMCELL Technologies, cat# 87701) and cryopreserved in 90% RPMI with 10% DMSO. Additional CD34+ BM mononuclear cells were purchased in thawed form from STEMCELL Technologies. Cryopreserved cells were thawed in 95% RPMI supplemented with 5% fetal calf serum (Gibco, cat# 26140079) and penicillin/streptomycin (Invitrogen).

**Cell culture for stimulation assay**

CD34+ BMMCs were isolated using the CD34 MicroBead Magnetic Separation Kit with MS Columns (Miltenyi Biotec, cat# 130-046-701). $CD34^{-}$ BMMCs and CD34+ cells were mixed to achieve 50% CD34+ enrichment and distributed into four wells in RPMI with penicillin/streptomycin and 5% FBS. Cytokines supporting B-cell development and candidate regulatory factors were added per Supporting Information S2: Table 3. Cells were incubated at 37°C in 5% $CO_2$ for 24 hours, then harvested for ScaleBio scRNA-seq.

***In Vitro* B-cell development co-culture for BDNF validation**

A six-well plate was pre-coated with an ~80% confluent HS-5 stromal cell layer. CD34+ BMMCs ($50\times10^{3}$ cells per well) were co-cultured over the HS-5 layer as previously described, maintained at 37°C in 5% $CO_2$ for three weeks, with media and cytokines refreshed twice weekly. Cells were harvested at specified time points, pelleted at 300×g for 5 minutes, and immunophenotyped by flow cytometry using the staining panel and gating strategy described in Supporting Information S1: Figure 4 and Supporting Information S2: Table 4. Events were analyzed using Cytobank software.

**Supplemental Figures**

**Supporting information, Figure 1: The expression levels of known B-cell markers correlates with biological expectations**

Violin plots showing the distribution of normalized expression levels of known B-cell markers: CD34, MME, CD19, MS4A1, CD79A and TNFRSF17 across cell clusters. Dot plots within each violin represent individual cell expression values derived from GSE144938 data (**A**) and GSE120446 (**B**). CD34 is highly expressed by HSPCs and the subsequent markers: MME (CD10), CD19, MS4A1 (CD20) and CD79A progressively appear until the expression of TNFRSF17 (BCMA) that characterizes plasma cells.

**Supporting information, Figure 2: Differentially expressed B-cell genes have different Waldstat values**

Reverse cumulative distribution of Wald statistics across genes. The plot shows the number of genes (y-axis) with a Wald statistic greater than or equal to a given threshold (x-axis). Each point along the curve represents the cumulative count of genes that exceed the corresponding WaldStat value. The red dashed vertical line indicates a threshold of WaldStat = 120, above which 1,556 genes remain.

**Supporting information, Figure 3: Candidates exhibit distinct specificities for non-B-cell populations in the bone marrow**

We tested the binding of each of the 67 candidates to receptors expressed on 11 distinct non-B-cell subpopulations within the BM: BNK, cMOP, CMP, GMP, MDP, MEP, MLP, MPP, NKP, preM and proN. A heatmap showing the putative interactions between the candidates (Y-axis) and non-B-cell populations (X-axis) is shown. Green represents positive interactions i.e. the non-B-cell subpopulation significantly expresses one of the receptors that pair with the candidate. Orange indicates that no candidate receptor is significantly expressed by the non-B-cell subpopulation (avg_log2FC > 1, p_adj < 0.01). Candidates are ranked based on the number of populations they interact with. BNK- B NK progenitors, cMOP- Common Monocyte Progenitor, CMP- Common Myeloid Progenitor, GMP- Granulocyte-Monocyte Progenitor, MDP- Monocyte-Derived Macrophages, MEP- Megakaryocyte-Erythroid Progenitor, Lymphoid Primed Multipotent Progenitor, MPP- Multipotent Progenitor, NKP- Natural Killer Progenitor, preM- Pre-monocyte, proN- Pro-Neutrophil.

**Supporting information, Figure S4: Gating strategy for characterizing B cells in flow cytometry** Flow cytometry scatter plots depict representative gating strategies for exploring the development of B cells. After gating on single cells, DAPI cells (viable cells) were selected and tested for expression of CD34, CD10 and CD19 markers.

**Supporting information, Figure S5: CXCL12 and TGFB3 upregulated and downregulated immune and developmental related pathways respectively**

Dot plots of GSEA results illustrate biological processes associated with immune and developmental pathways following simulations with CXCL12 **(A)** and TGFB3 **(B)**. Gene count refers to the number of genes associated with each REACTOME biological process. Gene ratio is the percentage of genes that are significantly correlated with the REACTOME biological process from the total number of genes associated with that process. Terms are ranked in the figure by decreasing gene ratio.

**Supporting information, Figure 6: BDNF acts at the beginning of the trajectory**

**(A)** Scaled expression of the BDNF receptor, NGFRAP1, along the B-cell developmental trajectory,  as derived from scRNA-seq data of healthy human BM samples (GSE149938) ( Methods). Each dot represents a cell. Low and high expressions of NGFRAP1 are colored as blue and red, respectively.

(**B**) An associated graph displays the scaled expression pattern along the trajectory.

**Supporting information, Figure 7: An *in-vitro* biological system supports the development of B cells**

An *in-vitro* biological system, where isolated $CD34^+$ cells from healthy human bone marrow samples were co-cultured with MSCs. The cells were analyzed by flow cytometry once a week for 21 days.

**Supporting information, Figure 8: The response pattern of putative regulators following Rituximab treatment is shared among those whose receptors behave similarly across the development of B cells**

An illustration showing the upregulation/downregulation of putative regulators acting at the end/beginning of the trajectory accordingly, following induction of B lymphopoiesis by Rituximab treatment. Rituximab destroys $CD20^{+}$ progenitor B cells, therefore less “late” receptors are available to bind their ligands leading to their increased levels. The compensated proliferation of early $CD20^{-}$ progenitor B cells, leads to an increased level of “early” receptors and lower levels of their ligands.

**Supporting information, Figure 1: The expression levels of known B-cell markers correlates with biological expectations**

**A**

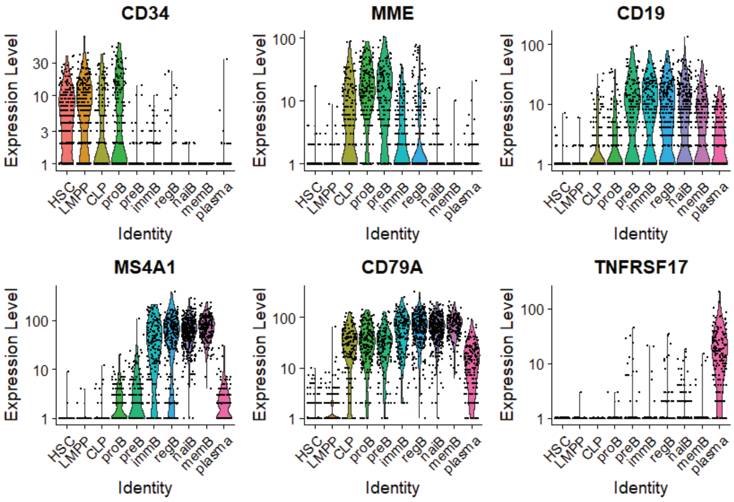


**B**

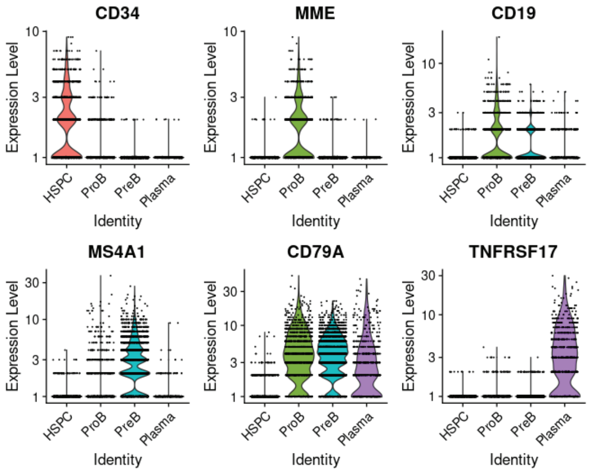

**Supporting information, Figure 2: Differentially expressed B-cell genes have different Waldstat values**

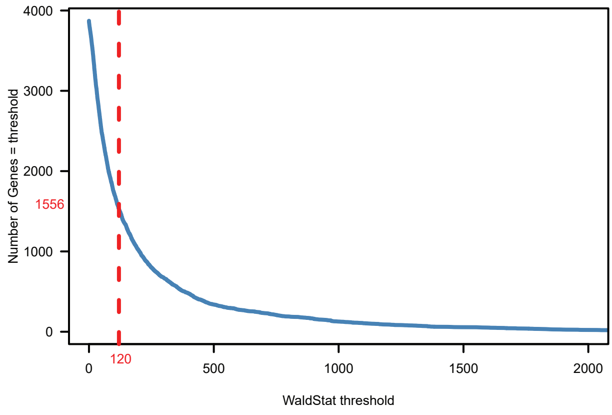

### Supporting information, Figure 3: Candidates exhibit distinct specificities for non-B- cell populations in the bone marrow

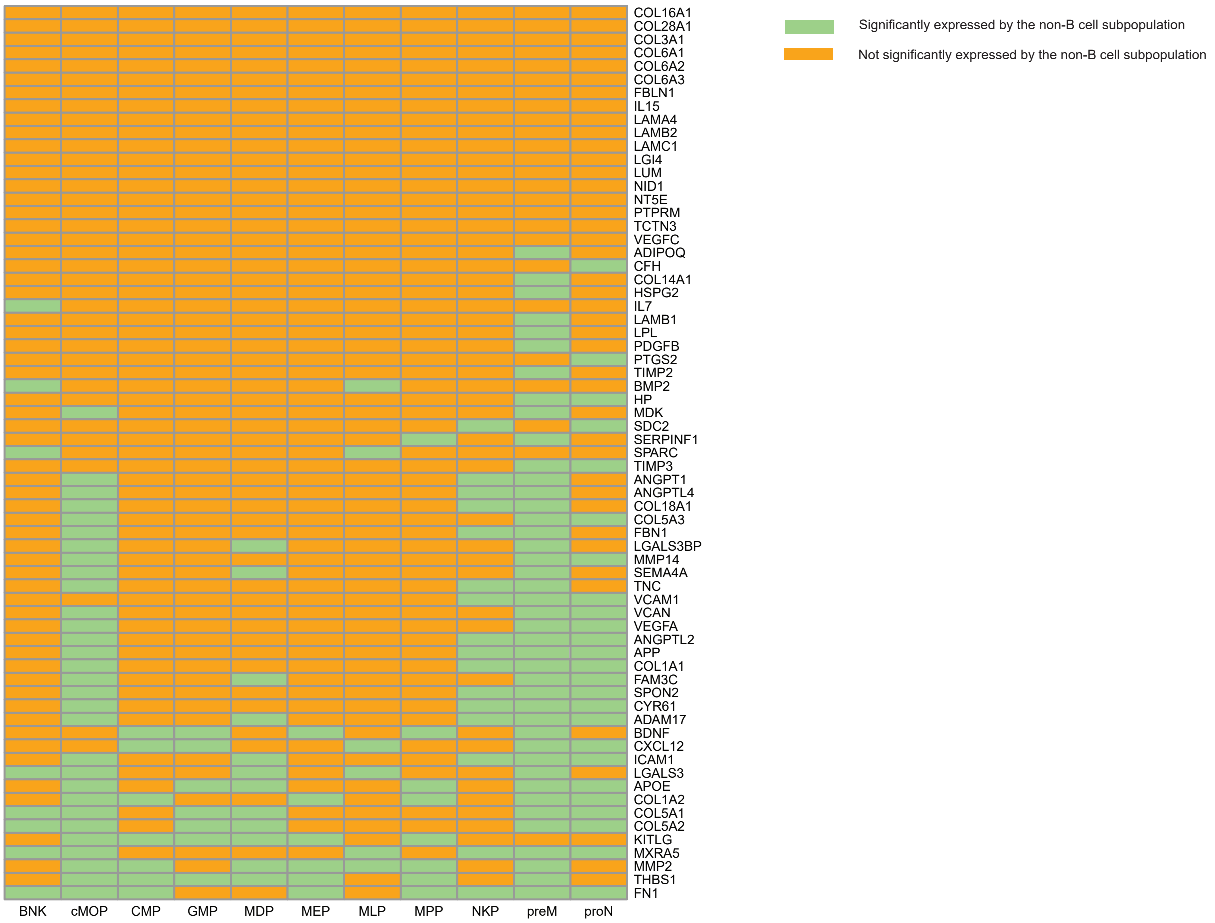

**Supporting information, Figure S4: Gating strategy for characterizing B-cells in flow cytometry**

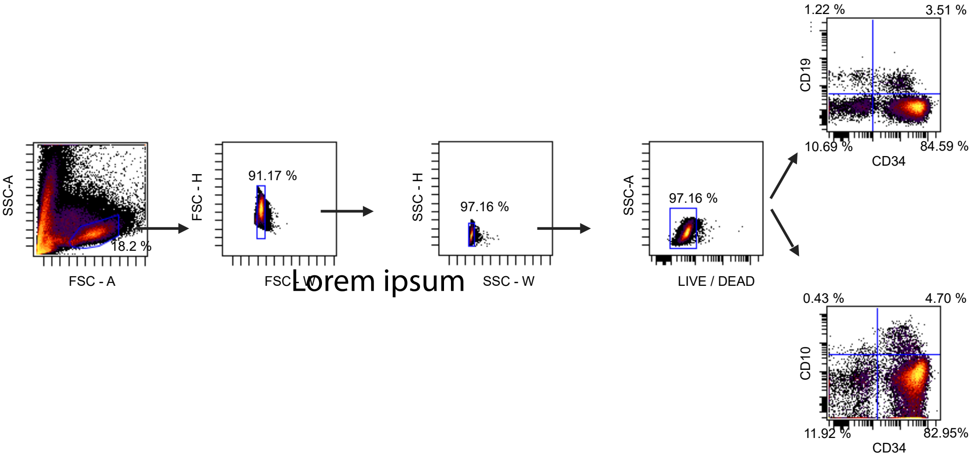

**Supporting information, Figure S5: CXCL12 and TGFB3 upregulated and downregulated immune and developmental related pathways respectively**

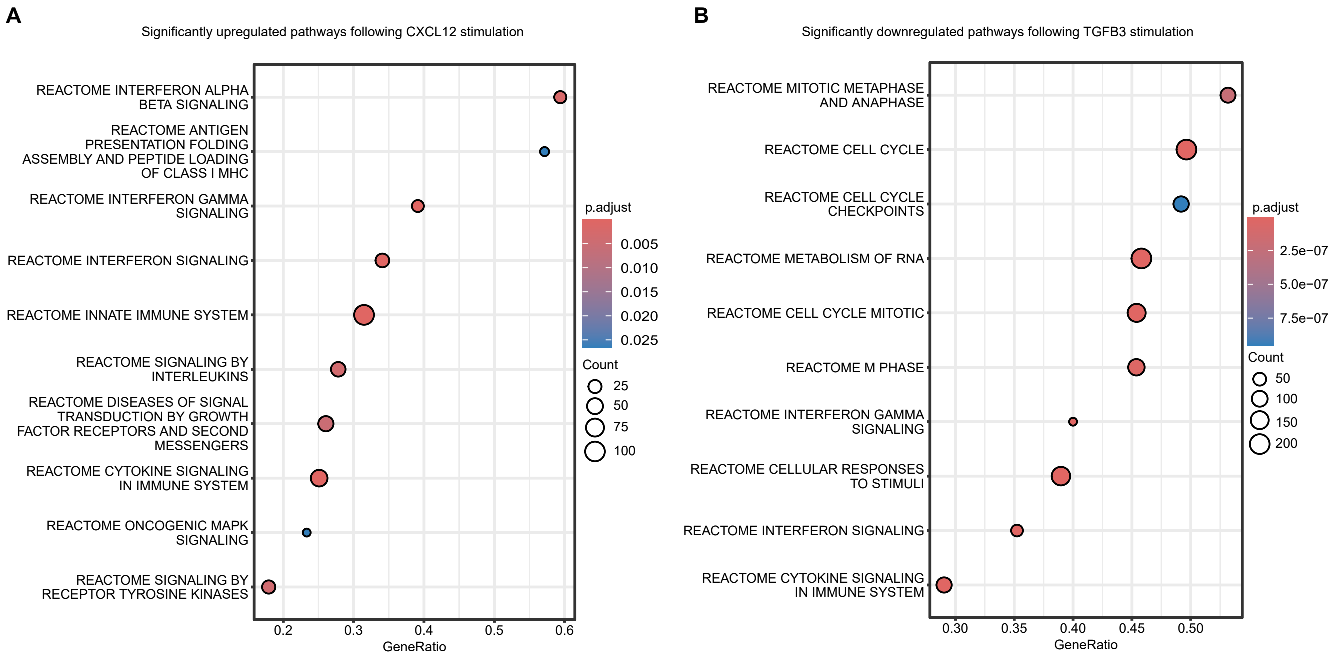

**Supporting information, Figure 6: BDNF acts at the beginning of the trajectory**

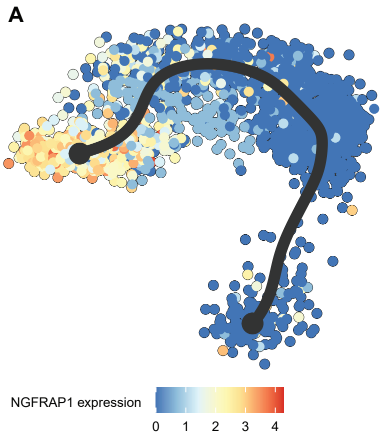


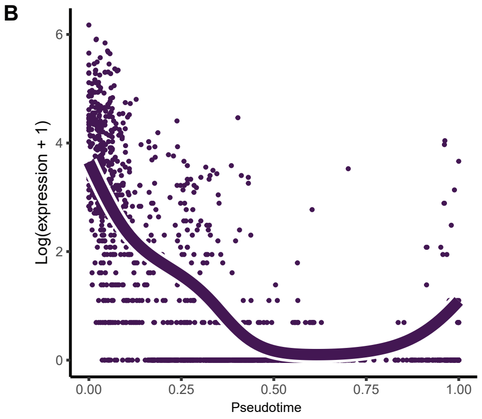

**Supporting information, Figure 7: An in-vitro biological system supports the development of B-cells**

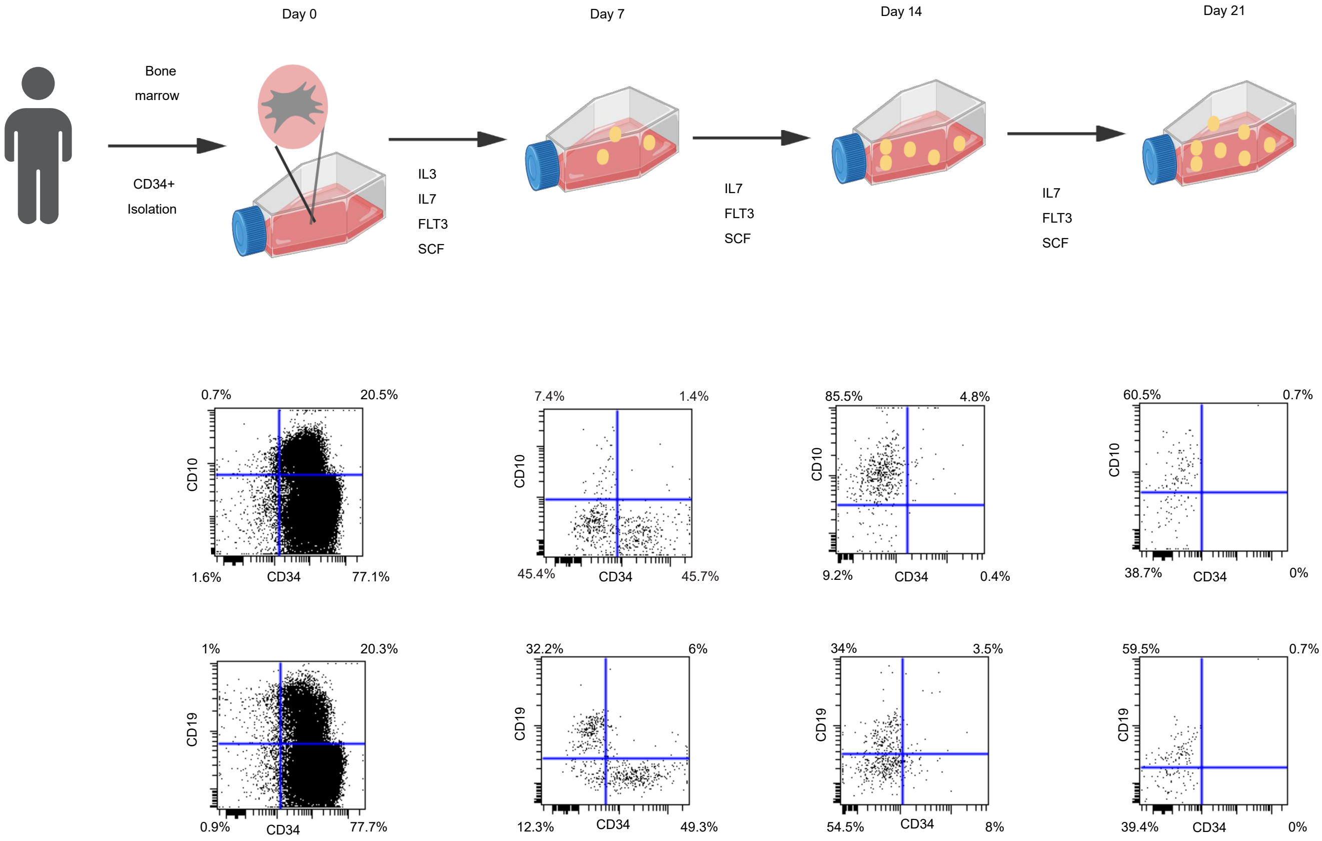

**Supporting information, Figure 8: The response pattern of putative regulators following Rituximab treatment is shared among those whose receptors behave similarly across the development of B-cells**

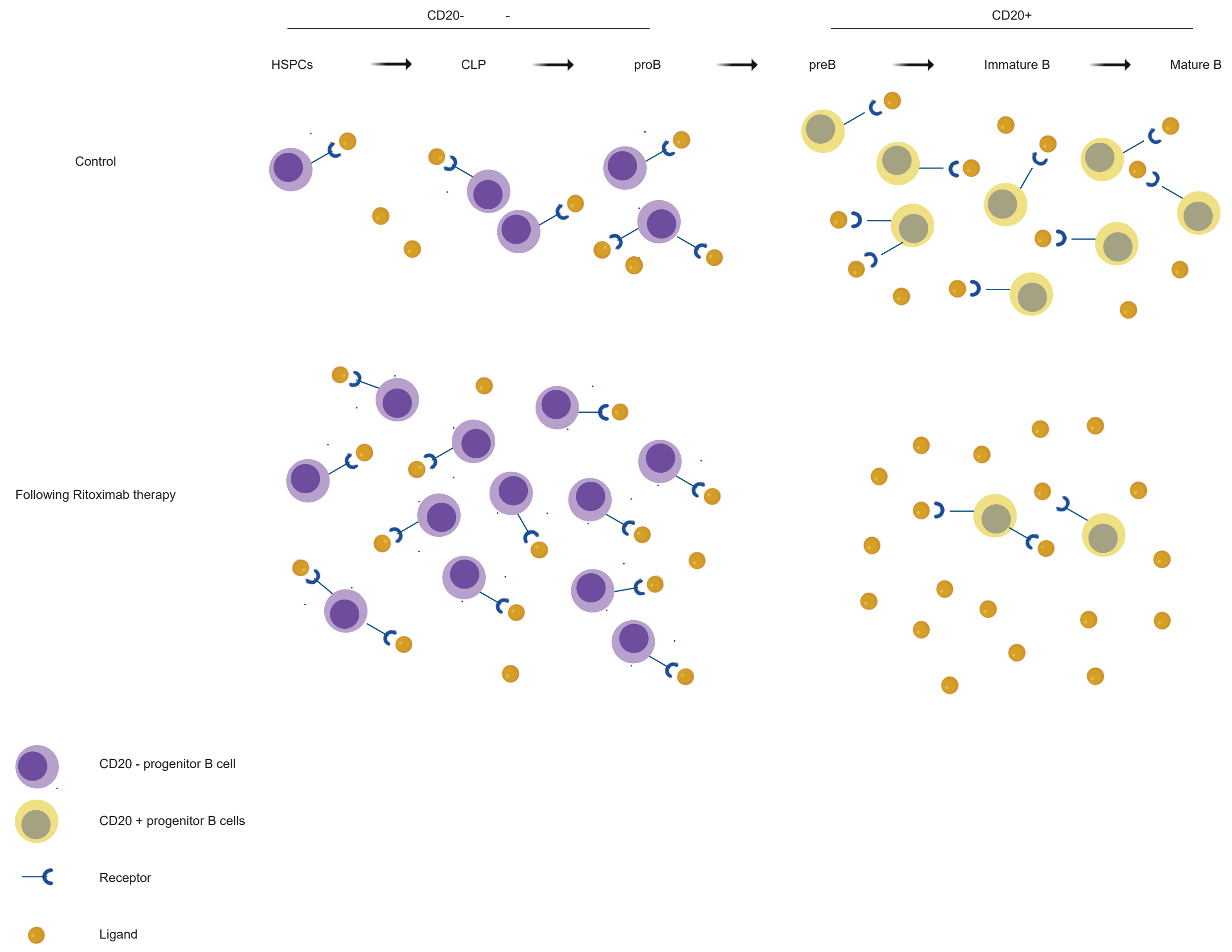
