## Supplementary figures and images for "A systems approach to B-cell development identifies BDNF as a regulator of human B lymphopoiesis"

### Graphical abstract

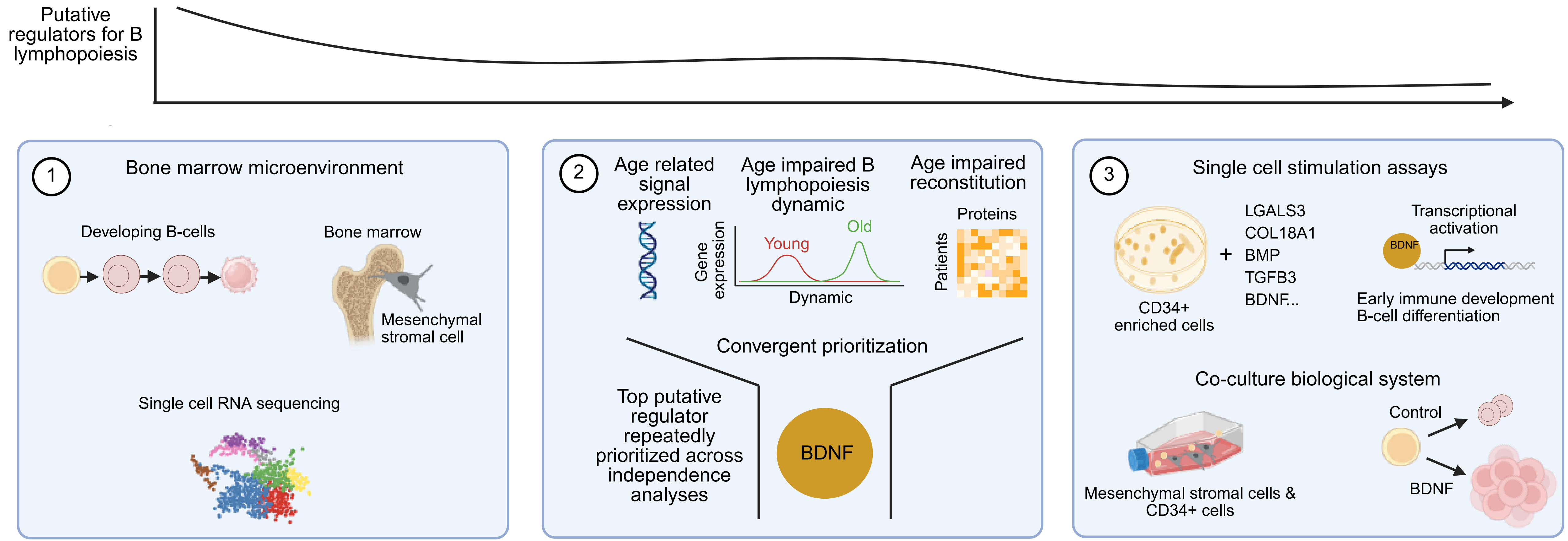


Potential strategy to improve immune reconstitution after prolonged B-cell depletion
